# Proximate, Mineral and Anti-nutrient Characteristics of Fermented Cassava Product (Pupuru) Supplemented with Fermented *Moringa oleifera* Seed

**DOI:** 10.64898/2026.10.13.758933

**Authors:** Gift Rebecca Asefon, Foluso Olutope Adetuyi, Kayode Olayele Karigidi

**Affiliations:** Department of Chemical Sciences, Olusegun Agagu University of Science and Technology, Okitipupa, Ondo State, Nigeria; Department of Cell and Molecular Biology, University of Rhode Island, Kingston, RI, USA

**Keywords:** Pupuru, cassava, *Manihot esculenta*, *Moringa oleifera*, fermentation, proximate composition, mineral profile, anti-nutrients, food fortification, Nigeria

## Abstract

**Background:** Pupuru, a traditionally fermented and smoked-dried cassava (*Manihot esculenta*) product, is widely consumed in South-Western Nigeria but is limited by low protein content and poor nutritional profile.

**Objective:** This study investigated the proximate composition, mineral profiles, and anti-nutritional factors of Pupuru supplemented with fermented *Moringa oleifera* seeds at varying inclusion levels.

**Methods:** Fermented cassava mash was supplemented with fermented *Moringa oleifera* seeds at 0% (control), 1%, 5%, and 10% inclusion levels. The resulting flour blends were analyzed for proximate composition, macro- and micromineral content, and anti-nutritional factors using standard AOAC methods.

**Results:** Supplementation significantly (p<0.05) increased protein (1.43–4.33%), fat (0.28–0.87%), fiber (0.32–0.75%), ash (0.40–1.30%), and moisture (12.57–13.06%) contents while carbohydrate content decreased (84.52–80.20%). Mineral profiles improved significantly across all treatment levels for iron (18.28–22.40 mg/kg), zinc (7.71–12.51 mg/kg), manganese (3.66–8.91 mg/kg), and copper (0.67–1.14 mg/kg). Anti-nutritional factors including phytate (0.27–0.30%), saponin (0.05–0.09%), oxalate (0.13–0.16%), and tannin remained within acceptable limits.

**Conclusion:** *Moringa oleifera* seed supplementation significantly improved the nutritional quality of Pupuru without compromising its safety profile, presenting a viable strategy for addressing protein-calorie malnutrition and mineral deficiencies in cassava-consuming populations.

## 1. Introduction

Cassava (*Manihot esculenta* Crantz) ranks as the third most important source of dietary energy after rice and maize for human populations and livestock across the world, particularly in sub-Saharan Africa (Abotbina et al., 2022). Its adaptability to marginal soils, drought tolerance, and year-round availability make it a critical food security crop for millions of smallholder farmers in developing countries (Adjei, 2021; Bayata, 2019). In Nigeria alone, cassava is processed into numerous traditional food products that serve as dietary staples for diverse population groups.

Among the various cassava-derived products, Pupuru occupies a unique cultural and nutritional position. Pupuru is a traditionally fermented, smoked-dried cassava food native to the Ilaje and Ikale tribes of Ondo State and widely consumed in the riverine communities of South-Western, Eastern, and Middle Belt regions of Nigeria (Isaac-Bamgboye et al., 2020). The product is consumed by an estimated 4–6 million people in Nigeria and across several other African nations (Adejuyitan et al., 2018). Despite its widespread consumption, Pupuru is recognized as an energy-dense but nutritionally limited food, primarily providing carbohydrates while remaining significantly deficient in protein, essential minerals, and vitamins (Obafemi et al., 2022; Akinjayeju et al., 2019).

Protein-energy malnutrition (PEM) and micronutrient deficiencies remain persistent public health challenges across cassava-dependent communities in sub-Saharan Africa. The heavy reliance on cassava products such as Pupuru as dietary staples, without adequate complementary protein sources, contributes significantly to these nutritional deficits, particularly among vulnerable groups including children, pregnant women, and women of reproductive age (Animashahun et al., 2022). Strategies to enhance the nutritional value of cassava-based foods through fortification and supplementation have therefore attracted considerable scientific interest.

*Moringa oleifera* Lam., commonly known as the drumstick tree or miracle tree, has emerged as a promising nutritional supplement owing to its exceptional nutrient density. The seeds, leaves, and pods of *M. oleifera* are rich sources of protein, essential minerals including calcium, potassium, iron, and phosphorus, as well as vitamins and bioactive phytochemicals (Gopalakrishnan et al., 2016; Matic et al., 2018). Notably, *M. oleifera* seeds contain appreciable quantities of lipids, primarily oleic, stearic, and palmitic acids, alongside high-quality protein with a favorable amino acid profile (de la Mora-López et al., 2018). The plant is widely accessible in tropical and subtropical regions of Africa, making it a cost-effective and sustainable nutritional supplement for rural communities.

Fermentation as a processing technique has been demonstrated to improve the bioavailability of nutrients, reduce anti-nutritional factors, and enhance the sensory properties of food products (Halake and Chinthapalli, 2020; Obafemi et al., 2022). The fermentation of both cassava and *Moringa oleifera* seeds prior to supplementation may therefore synergistically improve the nutritional profile of Pupuru while simultaneously reducing anti-nutrient concentrations that could otherwise limit mineral bioavailability.

Despite the nutritional potential of *M. oleifera* seeds and the widespread consumption of Pupuru, limited scientific information exists on the combined effect of fermented *M. oleifera* seed supplementation on the proximate composition, mineral profile, and anti-nutritional characteristics of Pupuru. This study therefore aimed to determine the proximate, mineral, and anti-nutrient characteristics of *Manihot esculenta* product (Pupuru) supplemented with fermented *Moringa oleifera* seeds at varying inclusion levels, with a view to providing scientific evidence for the use of *M. oleifera* seeds as a nutritional enhancer for cassava-based staple foods.

## 2. Materials and Methods

### 2.1 Plant Materials and Study Location

Matured cassava roots (*Manihot esculenta*) were purchased from Igodan market, Okitipupa, Ondo State, Nigeria, while matured *Moringa oleifera* seeds were sourced from Ekiti market, Ekiti State, Nigeria. Sample preparation and analyses were conducted at the Department of Chemical Sciences, Olusegun Agagu University of Science and Technology (OAUSTECH), Okitipupa, Ondo State, Nigeria.

### 2.2 Preparation of Fermented Cassava (Pupuru)

Cassava tubers were manually peeled, washed, and sliced into uniform pieces to ensure regular shape and size for consistent fermentation. The sliced cassava was weighed into four portions: 5.0 kg (control), 4.95 kg, 4.75 kg, and 4.50 kg, and each portion was separately submerged in water inside plastic containers. Spontaneous fermentation was allowed to proceed for five days at ambient temperature (28–32°C). Following fermentation, excess water was drained, and fibers were manually removed from the fermented mash.

### 2.3 Preparation of Fermented Moringa oleifera Seeds

*Moringa oleifera* seeds were manually dehulled and sun-dried for five hours. The dried seeds were weighed and blended into powder. Three separate portions of 0.05 kg (1%), 0.25 kg (5%), and 0.50 kg (10%) were individually soaked in one liter of water inside plastic containers and fermented for 120 hours (five days) at ambient temperature (28–32°C).

### 2.4 Production of Supplemented Pupuru Flour

Fermented *M. oleifera* seeds were incorporated into the fermented cassava mash at the following ratios (*Manihot esculenta*: *Moringa oleifera*, kg/kg): T1 — 5.0:0 (control); T2 — 4.95:0.05 (1%); T3 — 4.75:0.25 (5%); T4 — 4.50:0.50 (10%). The blended mashes were packed into sacks and subjected to mechanical pressing to remove excess water. The dewatered mash was sun-dried on sack mats, fried in aluminum pans without burning, and subsequently milled into fine Pupuru flour. Flour samples were packaged in sealed plastic bags pending analysis.

### 2.5 Proximate Analysis

Proximate composition was determined according to the official methods of the Association of Official Analytical Chemists (AOAC, 2005, 18th Edition). Crude protein was determined by the semi-micro Kjeldahl method (AOAC 988.05) using a conversion factor of 6.25. Crude fat was determined by Soxhlet extraction using petroleum ether (AOAC 2003.06). Moisture content was determined by oven-drying at 100°C for 24 hours (AOAC 967.08). Ash content was determined by incineration in a muffle furnace at 550°C for four hours (AOAC 942.05). Crude fiber was determined by sequential acid-alkaline digestion (AOAC 958.06). Nitrogen-free extract (carbohydrate) was calculated by difference: NFE = 100 – (% moisture + % crude protein + % crude fat + % crude fiber + % ash).

### 2.6 Mineral Analysis

Macro-minerals (calcium, potassium, sodium, phosphorus, and magnesium) and micro-minerals (iron, zinc, copper, manganese, and selenium) were determined following AOAC (975.11) methods. Ash digests were prepared using 2M HCl. Calcium, potassium, and sodium were determined using a Jenway Digital Flame Photometer (PFP7 Model). Phosphorus was determined spectrophotometrically using the vanadate-molybdate method at 470 nm (AOAC 975.16). Micro-mineral elements (selenium, magnesium, copper, manganese, iron, zinc) were determined using a Buck 200 Atomic Absorption Spectrophotometer (Buck Scientific Model 211vgp) at their respective wavelengths (AOAC 975.23).

### 2.7 Anti-nutritional Analysis

Phytic acid content was determined by the method of Young and Greaves (1940) as described by Reddy and Kove (1999), involving extraction with 2% HCl followed by titration with ferric chloride solution. Oxalate content was determined by the method of Day and Underwood (1986) using permanganate titration. Saponin content was determined by the gravimetric method of Obadoni and Ochuko (2001) using aqueous ethanol extraction and n-butanol purification. Alkaloid content was determined by the method of Harborne (1993). Tannin content was determined spectrophotometrically at 760 nm using ferric chloride reagent following the procedure of Evans and Trease (1989).

### 2.8 Statistical Analysis

All analyses were performed in triplicate and results expressed as mean ± standard deviation. Data were subjected to one-way analysis of variance (ANOVA) using SPSS version 23.0 (IBM Corp., Armonk, NY, USA). Means were separated using Duncan’s Multiple Range Test (DMRT) at p<0.05 significance level.

## 3. Results

### 3.1 Proximate Composition

The proximate composition of Pupuru supplemented with fermented *Moringa oleifera* seeds is presented in Table 1. Supplementation resulted in significant (p<0.05) increases in crude protein, crude fat, crude fiber, ash, and moisture contents, while carbohydrate content decreased progressively with increasing *M. oleifera* seed inclusion. Crude protein ranged from 1.43±0.06% in the control (T1) to 4.33±0.06% at the 10% supplementation level (T4). Crude fat increased from 0.28±0.01% (T1) to 0.87±0.02% (T4), while crude fiber increased from 0.32±0.01% (T1) to 0.75±0.01% (T4). Ash content ranged from 0.40±0.01% (T1) to 1.30±0.02% (T4), and moisture content ranged from 12.57±0.02% (T1) to 13.06±0.02% (T4). Carbohydrate content decreased significantly from 84.52±0.04% in the control to 80.20±0.12% at the highest supplementation level.

**Table 1.**
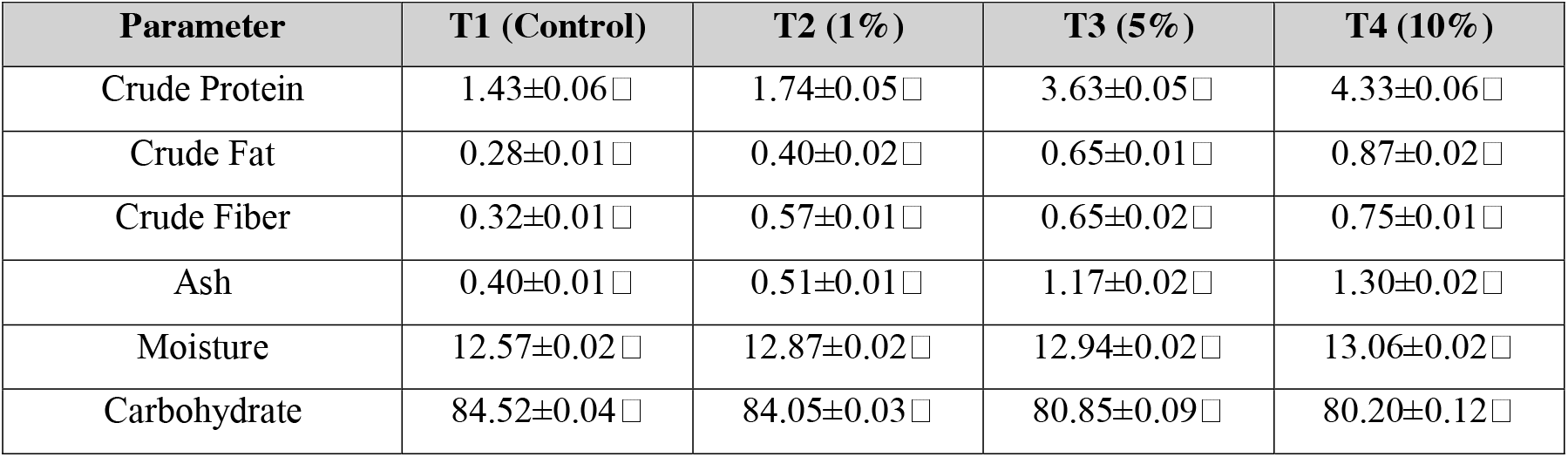
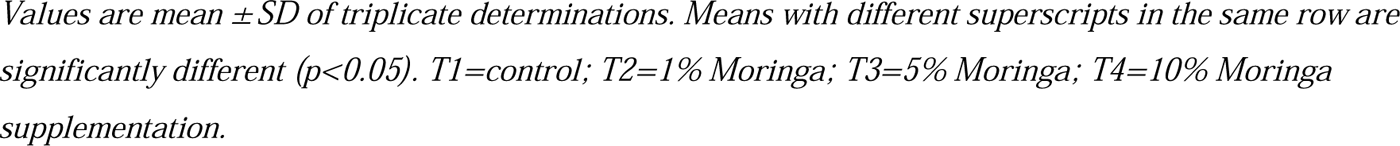
Proximate composition (%) of Pupuru supplemented with fermented Moringa oleifera seeds.

| Parameter | T1 (Control) | T2 (1%) | T3 (5%) | T4 (10%) |
| --- | --- | --- | --- | --- |
| Crude Protein | $1.43 \pm 0.06\%$ | $1.74 \pm 0.05\%$ | $3.63 \pm 0.05\%$ | $4.33 \pm 0.06\%$ |
| Crude Fat | $0.28 \pm 0.01\%$ | $0.40 \pm 0.02\%$ | $0.65 \pm 0.01\%$ | $0.87 \pm 0.02\%$ |
| Crude Fiber | $0.32 \pm 0.01\%$ | $0.57 \pm 0.01\%$ | $0.65 \pm 0.02\%$ | $0.75 \pm 0.01\%$ |
| Ash | $0.40 \pm 0.01\%$ | $0.51 \pm 0.01\%$ | $1.17 \pm 0.02\%$ | $1.30 \pm 0.02\%$ |
| Moisture | $12.57 \pm 0.02\%$ | $12.87 \pm 0.02\%$ | $12.94 \pm 0.02\%$ | $13.06 \pm 0.02\%$ |
| Carbohydrate | $84.52 \pm 0.04\%$ | $84.05 \pm 0.03\%$ | $80.85 \pm 0.09\%$ | $80.20 \pm 0.12\%$ |

Values are mean ± SD of triplicate determinations. Means with different superscripts in the same row are significantly different ( $p < 0.05$ ). T1=control; T2=1% Moringa; T3=5% Moringa; T4=10% Moringa supplementation.

### 3.2 Macromineral Composition

Table 2 presents the macromineral composition of the supplemented Pupuru flour blends. All macromineral values increased with increasing *M. oleifera* seed inclusion. Sodium ranged from 0.04±0.00% (T1) to 0.06±0.00% (T4), potassium from 0.12±0.00% to 0.15±0.00%, and calcium from 0.03±0.00% to 0.06±0.00%. Phosphorus increased significantly from 0.13±0.00% (T1) to 0.16±0.00% (T4), while magnesium increased from 0.04±0.00% (T1) to 0.07±0.00% (T4).

**Table 2.** Macromineral composition (%) of Pupuru supplemented with fermented Moringa oleifera seeds.

| Mineral | T1 (Control) | T2 (1%) | T3 (5%) | T4 (10%) |
| --- | --- | --- | --- | --- |
| Sodium | $0.04 \pm 0.00^a$ | $0.05 \pm 0.00^{ab}$ | $0.05 \pm 0.00^{ab}$ | $0.06 \pm 0.00^b$ |
| Potassium | $0.12 \pm 0.00^a$ | $0.13 \pm 0.00^a$ | $0.14 \pm 0.00^a$ | $0.15 \pm 0.00^a$ |
| Calcium | $0.03 \pm 0.00^a$ | $0.04 \pm 0.00^a$ | $0.04 \pm 0.00^a$ | $0.06 \pm 0.00^b$ |
| Phosphorus | $0.13 \pm 0.00^a$ | $0.14 \pm 0.00^a$ | $0.15 \pm 0.00^{ab}$ | $0.16 \pm 0.00^b$ |
| Magnesium | $0.04 \pm 0.00^a$ | $0.06 \pm 0.00^b$ | $0.06 \pm 0.00^b$ | $0.07 \pm 0.00^b$ |

Values are mean ± SD of triplicate determinations. Means with different superscripts in the same row are significantly different ( $p < 0.05$ ).

### 3.3 Micromineral Composition

Micromineral concentrations in the supplemented Pupuru flour are presented in Table 3. All microminerals increased significantly (p<0.05) with increasing *M. oleifera* seed inclusion. Iron increased from 18.28±0.02 mg/kg (T1) to 22.40±0.01 mg/kg (T4), zinc from 7.71±0.02 to 12.51±0.01 mg/kg, copper from 0.67±0.01 to 1.14±0.02 mg/kg, and manganese from 3.66±0.01 to 8.91±0.02 mg/kg. Selenium showed a significant difference between the control and all supplemented treatments, ranging from 0.07±0.00 mg/kg (T1) to 0.10±0.00 mg/kg (T4).

**Table 3.** Micromineral composition (mg/kg) of Pupuru supplemented with fermented Moringa oleifera seeds.

| Mineral | T1 (Control) | T2 (1%) | T3 (5%) | T4 (10%) |
| --- | --- | --- | --- | --- |
| Iron | 18.28±0.02<sup>a</sup> | 19.79±0.01<sup>a</sup> | 20.66±0.02<sup>a</sup> | 22.40±0.01<sup>a</sup> |
| Zinc | 7.71±0.02<sup>a</sup> | 9.33±0.02<sup>a</sup> | 10.35±0.02<sup>a</sup> | 12.51±0.01<sup>a</sup> |
| Copper | 0.67±0.01<sup>a</sup> | 0.84±0.01<sup>a</sup> | 1.04±0.02<sup>a</sup> | 1.14±0.02<sup>a</sup> |
| Manganese | 3.66±0.01<sup>a</sup> | 5.33±0.01<sup>a</sup> | 6.30±0.02<sup>a</sup> | 8.91±0.02<sup>a</sup> |
| Selenium | 0.07±0.00<sup>a</sup> | 0.09±0.00<sup>a</sup> | 0.09±0.00<sup>a</sup> | 0.10±0.00<sup>a</sup> |

Values are mean ± SD of triplicate determinations. Means with different superscripts in the same row are significantly different ( $p<0.05$ ).

### 3.4 Anti-nutritional Composition

The anti-nutritional composition of the supplemented Pupuru is presented in Table 4. Saponin content was highest in the control (0.09±0.00%) and lowest in T2 (0.05±0.00%), with significant differences among treatments. Tannin content was negligible in T1 and T2 (0.00±0.00%) and very low in T3 and T4 (0.01±0.00%), with no significant difference between T3 and T4. Phytate content ranged from 0.27±0.00% to 0.30±0.00%, with the highest values in T2 and T3. Oxalate content decreased progressively from T2 (0.16±0.00%) to T4 (0.13±0.00%), with significant differences among all treatments.

**Table 4.** Anti-nutritional composition (%) of Pupuru supplemented with fermented Moringa oleifera seeds.

| Anti-nutrient | T1 (Control) | T2 (1%) | T3 (5%) | T4 (10%) |
| --- | --- | --- | --- | --- |
| Saponin | 0.09±0.00<sup>a</sup> | 0.05±0.00<sup>b</sup> | 0.08±0.00<sup>a</sup> | 0.07±0.00<sup>a</sup> |
| Tannin | 0.00±0.00<sup>a</sup> | 0.00±0.00<sup>a</sup> | 0.01±0.00<sup>a</sup> | 0.01±0.00<sup>a</sup> |
| Phytate | 0.29±0.00<sup>a</sup> | 0.30±0.00<sup>a</sup> | 0.30±0.00<sup>a</sup> | 0.27±0.00<sup>a</sup> |
| Oxalate | 0.15±0.00<sup>a</sup> | 0.16±0.00<sup>a</sup> | 0.14±0.00<sup>a</sup> | 0.13±0.00<sup>a</sup> |

Values are mean ± SD of triplicate determinations. Means with different superscripts in the same row are significantly different ( $p<0.05$ ).

## 4. Discussion

### 4.1 Proximate Composition

The progressive increase in crude protein content observed with increasing *M. oleifera* seed supplementation is consistent with the well-documented high protein content of *Moringa oleifera* seeds (Mgbemena and Obodo, 2016; Islam et al., 2021). The protein content of the 10% supplemented Pupuru (4.33%) represented a three-fold increase over the control (1.43%), which reflects the significant protein contribution of *M. oleifera* seeds. This finding aligns with observations reported by Oluwamukomi and Akinlabi (2011) for Pupuru supplemented with African yam bean seed flour. The enrichment of Pupuru protein content through *M. oleifera* supplementation holds considerable public health significance, as protein deficiency remains a critical nutritional challenge among populations dependent on cassava-based diets. The observed protein values, while improved, remain relatively low compared to recommended dietary requirements, suggesting that higher supplementation levels or combination with other high-protein ingredients may be warranted for populations at high risk of protein-energy malnutrition.

The significant increase in crude fat content with *M. oleifera* supplementation is attributable to the naturally high lipid content of *Moringa* seeds, which are predominantly composed of oleic, stearic, and palmitic acids (de la Mora-López et al., 2018; Nadeem and Imran, 2016). The increased fat content of supplemented Pupuru enhances its palatability and energy density, which is advantageous in food-insecure settings where energy intake maximization is desirable (Oluwamukomi and Akinlabi, 2011). The elevation in crude fiber content is nutritionally beneficial, as dietary fiber promotes gastrointestinal health, modulates glycemic response, and reduces the risk of non-communicable diseases including cardiovascular disease and type 2 diabetes.

The significant increase in ash content with progressive supplementation reflects the higher mineral content contributed by Moringa seeds and serves as an indirect indicator of improved mineral composition in the supplemented flour. The modest but significant increase in moisture content across treatments is within acceptable limits for food safety and shelf stability of dried flour products. The corresponding decrease in carbohydrate content is an expected consequence of ingredient dilution as protein-, fat-, and mineral-rich Moringa seeds replace a proportion of the carbohydrate-dense cassava base.

### 4.2 Mineral Composition

The significant enhancement of both macro- and micromineral profiles in supplemented Pupuru confirms previous reports of the high mineral density of *M. oleifera* seeds, particularly for potassium, sodium, phosphorus, and iron (Mgbemena and Obodo, 2016; Matic et al., 2018). The substantial increase in iron content from 18.28 mg/kg in the control to 22.40 mg/kg at 10% supplementation is of particular nutritional significance. Iron deficiency anemia remains the most prevalent micronutrient deficiency globally, disproportionately affecting women of reproductive age, pregnant women, and young children in sub-Saharan Africa (Animashahun et al., 2022). The enriched iron content of supplemented Pupuru suggests potential as a dietary strategy for addressing iron deficiency in vulnerable population groups.

The significant increases in zinc, copper, and manganese concentrations with *M. oleifera* supplementation are consistent with the established role of these trace elements in supporting immune function, enzymatic activity, antioxidant defense, and bone metabolism (Alozie et al., 2017). Zinc, in particular, plays critical roles in growth, wound healing, and immune competence, and its deficiency is prevalent in populations subsisting on cereal- and tuber-based diets with high phytate content. The enhancement of phosphorus and magnesium content in supplemented Pupuru is also nutritionally relevant, as these minerals regulate acid-base balance, support skeletal development, and facilitate energy metabolism (Alozie et al., 2017).

### 4.3 Anti-nutritional Composition

The low levels of anti-nutritional factors detected across all treatment groups reflect the effectiveness of combined fermentation and thermal processing (frying) in reducing anti-nutrient concentrations in both cassava and *Moringa oleifera* seeds (Ufere et al., 2021; Halake and Chinthapalli, 2020). Phytate concentrations across all treatments (0.27–0.30%) were substantially lower than the 624 mg/100g (approximately 0.62%) reported for raw cassava roots, confirming the efficacy of fermentation in reducing phytate through the activity of endogenous phytase and phosphatase enzymes (Taiwo, 2009; Soetan and Oyewole, 2009). Phytate binds divalent cations including iron, zinc, calcium, and magnesium, forming insoluble complexes that reduce mineral bioavailability (Sumayya et al., 2020). The low phytate levels observed therefore support the bioavailability of the enhanced mineral content in supplemented Pupuru.

The progressive reduction in oxalate content with higher Moringa supplementation levels (from 0.16% at 1% to 0.13% at 10%) is nutritionally favorable, as dietary oxalate at high concentrations can bind calcium and other minerals, reduce mineral absorption, and increase the risk of renal oxalate stone formation. The low tannin content across all treatments (0.00–0.01%) is consistent with the known tannin-reducing effects of fermentation and heat treatment. Tannin-rich foods impair protein digestibility, inhibit digestive enzyme activity, and reduce mineral absorption; therefore, the negligible tannin levels in supplemented Pupuru support its nutritional quality and digestibility.

The saponin content showed a non-linear pattern, with the control exhibiting the highest concentration (0.09%) and T2 the lowest (0.05%). While saponins at high concentrations can interfere with nutrient absorption, the low levels detected across all treatments are unlikely to pose significant anti-nutritional effects. Moreover, saponins at low concentrations possess documented biological activities including antioxidant, anti-inflammatory, and hypocholesterolemic properties (Soetan and Oyewole, 2009). Overall, the anti-nutritional factor profile of *M. oleifera*-supplemented Pupuru is well within safe limits for human consumption, supporting the safety and nutritional integrity of the product.

## 5. Conclusion

This study demonstrates that supplementation of Pupuru, a traditionally fermented cassava product, with fermented *Moringa oleifera* seeds significantly improves its proximate composition and mineral profile while maintaining anti-nutritional factors at safe and acceptable levels. The 10% supplementation level produced the most nutritionally enhanced product, yielding a three-fold increase in protein content, substantially higher micro- and macromineral concentrations, and improved fiber and fat contents relative to the unsupplemented control. These improvements, achieved through the combined action of fermentation and thermal processing, support the potential of fermented *M. oleifera* seeds as a cost-effective and locally available nutritional supplement for enhancing cassava-based staple foods.

The findings carry practical implications for addressing protein-energy malnutrition and micronutrient deficiencies — particularly iron, zinc, and protein — among cassava-dependent populations in sub-Saharan Africa. The use of *M. oleifera* seeds, which are widely available and underutilized across tropical Africa, represents a sustainable and culturally acceptable approach to food fortification that can be implemented at the household and community level. Future studies should investigate the sensory properties, functional characteristics, shelf stability, and in vivo mineral bioavailability of *M. oleifera*-supplemented Pupuru to further support its adoption as a nutritionally enhanced staple food.

## Declarations

### Author Contributions

G.R. Oluwasegun: Conceptualization, methodology, investigation, data collection, formal analysis, writing — original draft. F.O. Adetuyi: Supervision, project administration, writing — review and editing. K.O. Karigidi: Supervision, writing — review and editing.

### Conflicts of Interest

The authors declare no conflict of interest.

### Funding

This research received no external funding.

### Data Availability

The data supporting the findings of this study are available within the article.

## Notes

### Competing Interest Statement

The authors have declared no competing interest.

